# Very low depth whole genome sequencing in complex trait association studies

**DOI:** 10.1101/169789

**Authors:** Arthur Gilly, Lorraine Southam, Daniel Suveges, Karoline Kuchenbaecker, Rachel Moore, Giorgio E.M. Melloni, Konstantinos Hatzikotoulas, Aliki-Eleni Farmaki, Graham Ritchie, Jeremy Schwartzentruber, Petr Danecek, Britt Kilian, Martin O. Pollard, Xiangyu Ge, Emmanouil Tsafantakis, George Dedoussis, Eleftheria Zeggini

**Affiliations:** Department of Human Genetics, Wellcome Sanger Institute, Wellcome Genome Campus, Hinxton, Cambridge CB10 1HH, UK; Wellcome Centre for Human Genetics, University of Oxford, Oxford OX3 7BN, UK; Center for Genomic Science of IIT@SEMM, Fondazione Istituto Italiano di Tecnologia (IIT), Via Adamello 16, 20139, Milan, Italy; Department of Nutrition and Dietetics, School of Health Science and Education, Harokopio University of Athens, Greece; European Bioinformatics Institute, Wellcome Genome Campus, Hinxton CB10 1SH, UK; Anogia Medical Centre, Anogia, Greece

## Abstract

**Motivation:** Very low depth sequencing has been proposed as a cost-effective approach to capture low-frequency and rare variation in complex trait association studies. However, a full characterisation of the genotype quality and association power for very low depth sequencing designs is still lacking.

**Results:** We perform cohort-wide whole genome sequencing (WGS) at low depth in 1,239 individuals (990 at 1x depth and 249 at 4x depth) from an isolated population, and establish a robust pipeline for calling and imputing very low depth WGS genotypes from standard bioinformatics tools. Using genotyping chip, whole-exome sequencing (WES, 75x depth) and high-depth (22x) WGS data in the same samples, we examine in detail the sensitivity of this approach, and show that imputed 1x WGS recapitulates 95.2% of variants found by imputed GWAS with an average minor allele concordance of 97% for common and low-frequency variants. In our study, 1x further allowed the discovery of 140,844 true low-frequency variants with 73% genotype concordance when compared to high-depth WGS data. Finally, using association results for 57 quantitative traits, we show that very low depth WGS is an efficient alternative to imputed GWAS chip designs, allowing the discovery of up to twice as many true association signals than the classical imputed GWAS design.

**Supplementary Data:** Supplementary Data are appended to this manuscript.

## Introduction

The contribution of low-frequency and rare variants to the allelic architecture of complex traits remains largely unchartered. Power to detect association is central to genetic studies examining sequence variants across the full allele frequency spectrum. Whole genome sequencing (WGS)-based association studies hold the promise of probing a larger proportion of sequence variation compared to imputed genome-wide genotyping arrays. However, although large-scale high-depth WGS efforts are now underway (Brody, et al., 2017), comparatively high costs do not yet allow for the generalised transposition of the GWAS paradigm to high-depth sequencing. As sample size and haplotype diversity are more important than sequencing depth in determining power for association studies (Alex Buerkle and Gompert, 2013; Le and Durbin, 2011), low-depth WGS has emerged as an alternative, cost-efficient approach to capture low-frequency variation in large studies. Improvements in calling algorithms have enabled robust genotyping using WGS at low depth (4x-8x), leading to the creation of large haplotype reference panels (1000 Genomes Project Consortium, et al., 2015; McCarthy, et al., 2016), and to several low-depth WGS-based association studies (Astle, et al., 2016; Tachmazidou, et al., 2017; UK10K Consortium, et al., 2015). Very low depth (<2x) sequencing has been proposed as an efficient way to further improve the cost efficiency of sequencing-based association studies. Simulations have shown that in WES designs, extremely low sequencing depths (0.1-0.5x) are effective in capturing single-nucleotide variants (SNVs) in the common (MAF>5%) and low-frequency (MAF 1-5%) categories compared to imputed GWAS arrays (Pasaniuc, et al., 2012). The CONVERGE consortium demonstrated the feasibility of such approaches through the first successful case-control study of major depressive disorder in 4,509 cases and 5,337 controls (Converge Consortium, 2015), and we previously showed that 1x WGS allowed the discovery of replicating burdens of low-frequency and rare variants (Gilly, et al., 2016). However, a systematic examination of genotyping quality from 1x WGS and its implications for power in association studies is lacking, posing the question of the generalisability of such results in the wider context of next-generation association studies. Here, we perform very low depth (1x), cohort-wide WGS in an isolated population from Greece, show that imputation tools commonly used with chip data perform well using 1x WGS, and establish a detailed quality profile of called variants. We then demonstrate the advantages of 1x WGS compared to the more traditional imputed GWAS design both in terms of genotype accuracy and power to detect association signals.

## Results

As part of the Hellenic Isolated Cohorts (HELIC) study, we whole genome sequenced 990 individuals from the Minoan Isolates (HELIC-MANOLIS) cohort at 1x depth, on the Illumina HiSeq2000 platform. In addition, 249 samples from the MANOLIS cohort were sequenced at 4x depth (Southam, et al., 2017). Imputation-based genotype refinement was performed on the cohort-wide dataset using a combined reference panel of 10,244 haplotypes from MANOLIS 4x WGS, the 1000 Genomes (1000 Genomes Project Consortium, et al., 2015) and UK10K (UK10K Consortium, et al., 2015) projects (Figure 1).

**Figure 1:**
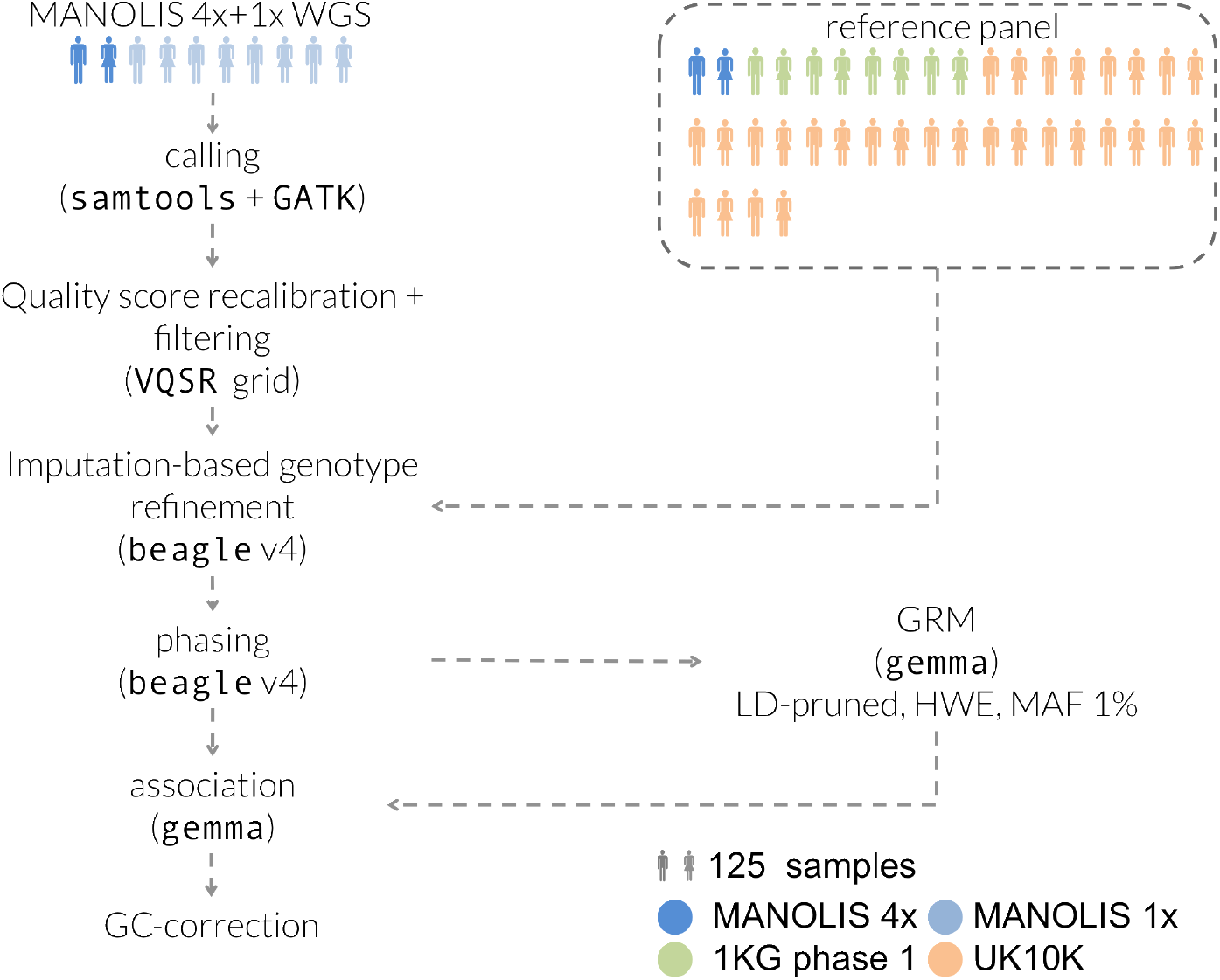
Processing pipeline for the MANOLIS 1x data. Tools and parameters for the genotype refinement and phasing steps were selected after benchmarking nine pipelines involving four different tools (See Methods).

### Variant calling pipeline

Prior to any imputation-based refinement, our approach allowed the capture of 80% and 100% of low-frequency (MAF 1-5%) and common (MAF>5%) SNVs, respectively, when compared to variants present on the Illumina OmniExpress and HumanExome chips genotyped in the same samples. In 10 control samples with high-depth WGS data downsampled to 1x, joint calling with MANOLIS resulted in pre-imputation false positive and false negative rates of 12% and 24.6%, respectively (See Methods).

In order to improve sensitivity and genotype accuracy, we compared nine genotype refinement and imputation pipelines using tools commonly used for genotyping chip imputation, using directly typed OmniExpress and ExomeChip genotypes as a benchmark (See Methods). We used a reference panel containing haplotypes from 4,873 cosmopolitan samples from the 1000 Genomes and UK10K projects, as well as the phased haplotypes from 249 MANOLIS samples sequenced at 4x depth. The best-performing pipeline, described in Figure 1, captures 95% of rare, 99.7% of low-frequency and 99.9% of common variants present in chip data, with an average minor allele concordance of 97% across the allele frequency spectrum (see Methods, Figure 2a., Supplementary Figure 1). 79.7% of 1x WGS variants were found using high-depth WGS at 22x in a subset of the MANOLIS samples (n=1,225), although this positive predictive value varied across the MAF spectrum, from 8.9% for singletons to 95.1% for common variants (Figure 2b.). Genotype concordance was similar, although slightly lower, when compared to the chip variants. Due to the 22x data being aligned to a different build, we were unable to compute genome-wide false positive rates, however by comparing 1x calls with those produced by whole-exome sequencing in 5 individuals from the MANOLIS cohort, we estimate a false-positive rate of 2.4% post-imputation in the coding parts of the genome (see Methods).

**Figure 2:**
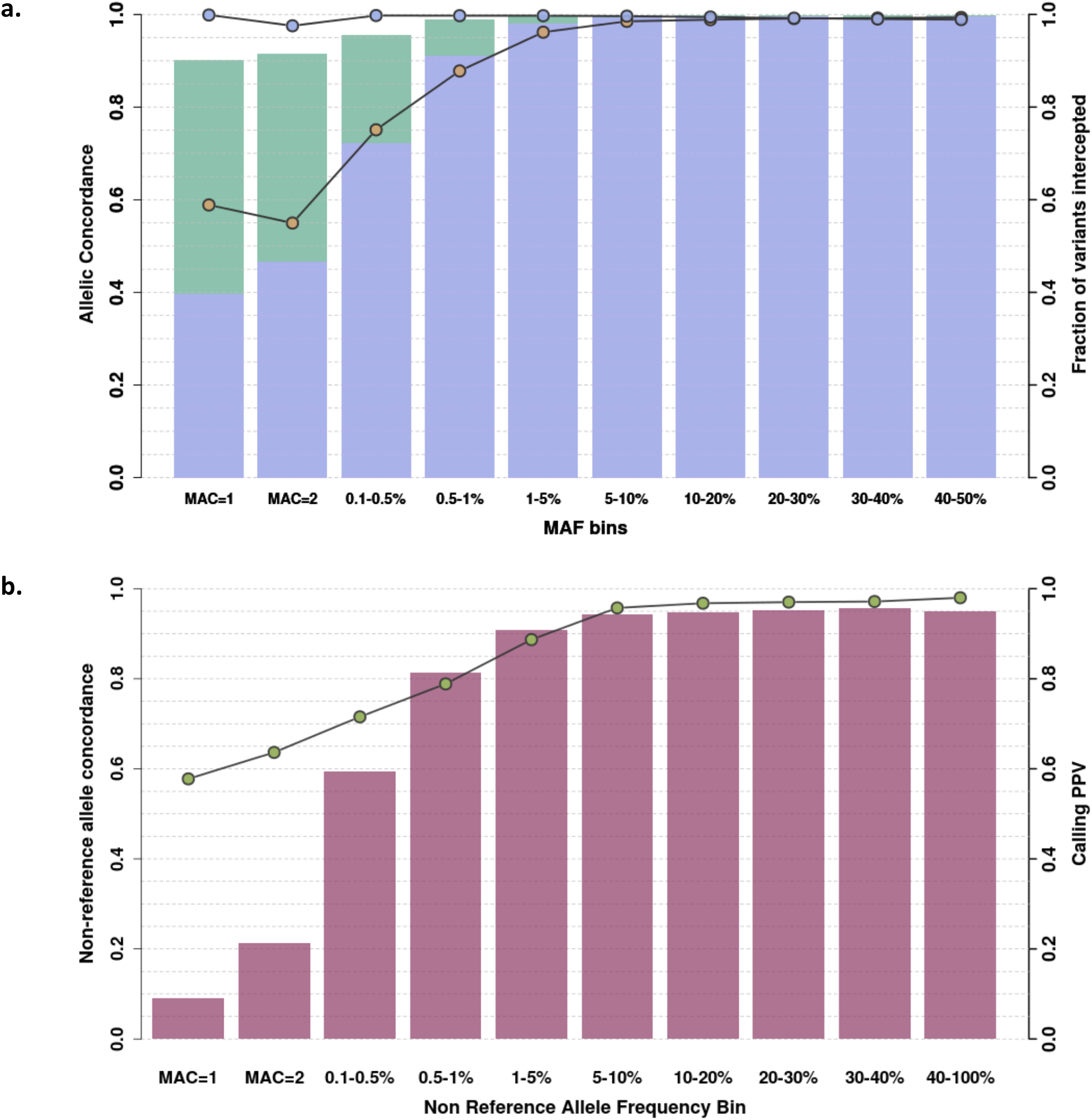
Concordance and call rate for very low depth WGS genotypes. **a**. Genotype (blue circles) and minor allele (yellow circles) concordance is computed for 1,239 samples in MANOLIS against merged OmniExpress and ExomeChip data. Call rate is assessed for the refined (purple) and refined plus imputed (green) datasets. **b**. Non-reference allele concordance (green circles) and positive predictive value (PPV) (fuchsia bars) is computed for 1,225 MANOLIS samples with both 22x WGS and low-depth calls.

### Comparison of variant call sets with an imputed GWAS

The genotype refinement and imputation step yielded 30,483,136 non-monomorphic SNVs in 1,239 MANOLIS individuals. The number of variants discovered using 1x WGS is nearly twice as high as that from array-based approaches. In a subset of 982 MANOLIS individuals with both 1x WGS, OmniExpress and ExomeChip data, we called 25,673,116 non-monomorphic SNVs using 1x WGS data, compared to 13,078,518 non-monomorphic SNVs in the same samples with chip data imputed up to the same panel (Southam, et al., 2017) without any imputation INFO score filtering. The main differences are among rare variants (MAF<1%) (Figure 3): 13,671,225 (53.2%) variants called in the refined 1x WGS are absent from the imputed GWAS, 98% of which are rare. 82% of these rare unique SNVs are singletons or doubletons, and therefore 9.5% of all variants called in the 1x WGS dataset were unique variants with MAC>2.

**Figure 3:**
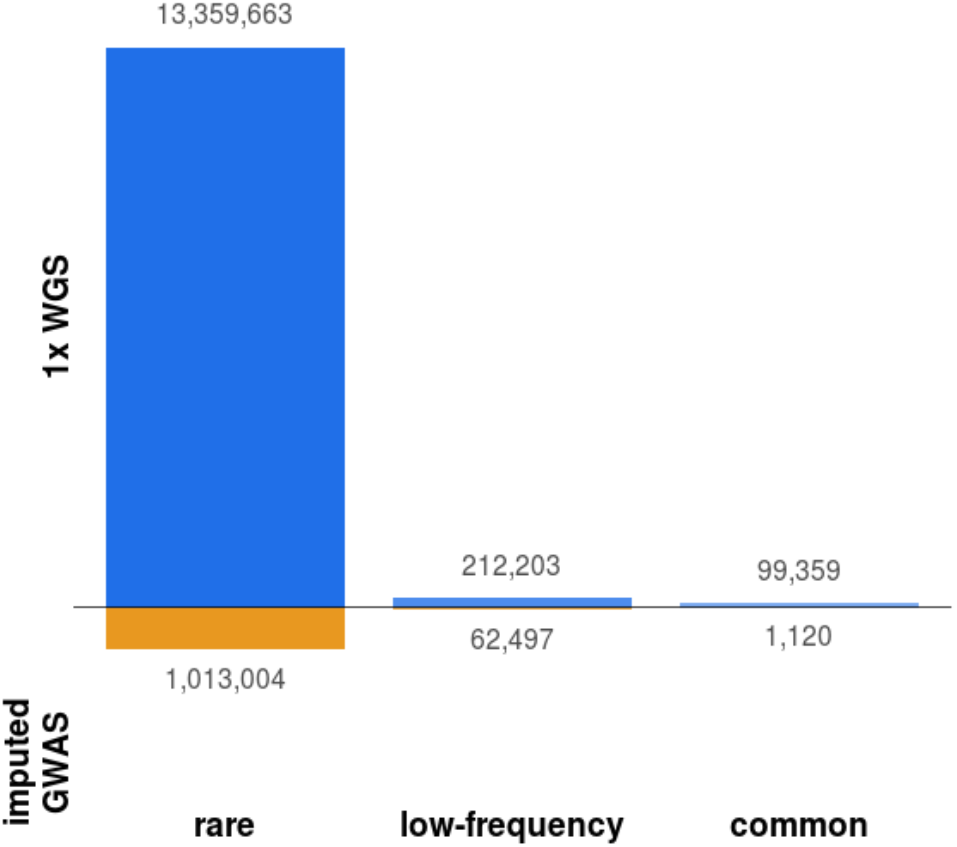
Unique variants called by sequencing and imputed GWAS. Variants unique to either dataset, arranged by MAF bin. Both datasets are unfiltered apart from monomorphics, which are excluded. MAF categories: rare (MAF<1%), low-frequency (MAF 1-5%), common (MAF>5%).

A crucial question is the proportion of true positives among these additional SNVs not found by GWAS and imputation. By comparing their positions and alleles with high-depth WGS in the same samples, we find that the PPV profile for these variants is much lower compared to when all variants are examined (Figure 4 and Figure 2.b). As expected, PPV is almost zero for additional singletons and doubletons, and just above 40% for the few additional common variants. 62% of low-frequency variants unique to the 1x are true positives, which corresponds to 140,844 low-frequency variants with high genotyping quality that are missed by the imputed GWAS. Minor allele concordance is lower than for all variants, with a lower bound at 55% for rare variants and reaching 73% for novel low-frequency variants.

**Figure 4:**
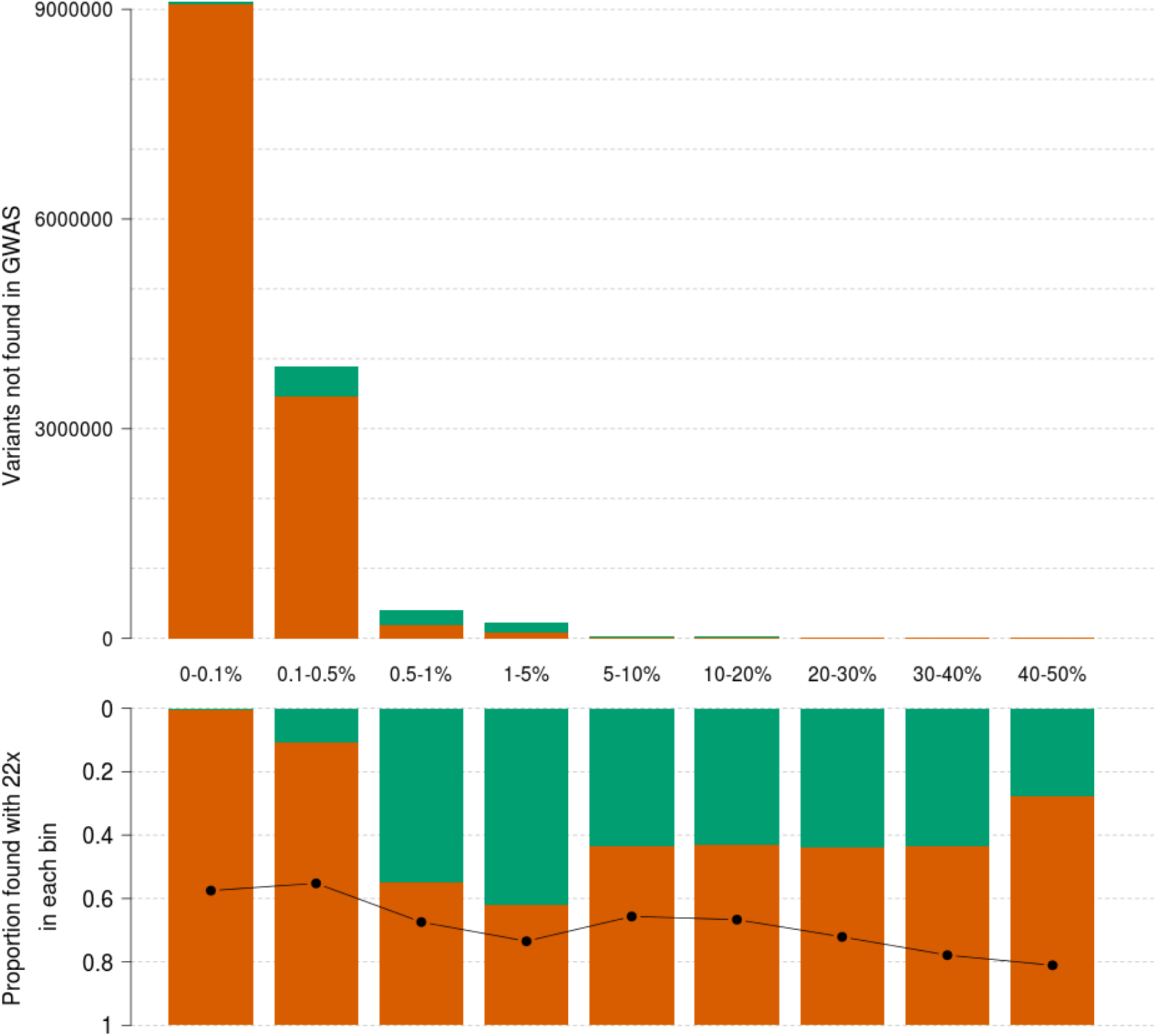
Positive predictive value of additional variants called in 1x sequencing. 1x variants not found in the GWAS data, arranged by MAF bin, in raw numbers (top). Green bars count variants recapitulated in the 22x (true positives). The proportion of these over the total (positive predictive value) is displayed in each bin in the bottom panel. The black line indicates minor allele concordance for true positive variants. The first category (0-0.1%) contains singletons and doubletons only.

### Comparison of association summary statistics with imputed GWAS

1x WGS calls a larger number of variants and is noisier than imputed GWAS in the same samples. To evaluate how this difference affects association study power, we performed genome-wide association of 57 quantitative traits in 1,225 overlapping samples with both imputed OmniExome and 1x WGS using both sources of genotype data. We then compared independent suggestively associated signals at p<5x10^−7^ (Supplementary Table 1). These signals were then cross-referenced with a larger (n=1,457) study based on 22x WGS on the same traits in the same cohort(Gilly, et al., 2018). We only considered signals to be true if they displayed evidence for association with at most a two order of magnitude attenuation compared to our suggestive significance threshold (P<5x10^−5^). According to this metric, 52 of 182 independent signals (28.5%) were true in the imputed GWAS, in contrast to 108 of 462 (23.4%) in the 1x study (Figure 5). With an equal sample size and identically transformed traits, 1x therefore allowed to discover twice as many independent GWAS signals with almost identical truth sensitivity. Seven rare and three suggestive low-frequency variant associations in the 1x WGS data (9.2% of all signals) were driven by a variant not present and without a tagging SNP at r^2^>0.8 in the imputed GWAS, whereas the converse is true for only two rare variants in the imputed GWAS. Among variants called or tagged in the imputed GWAS, 4 rare, 11 low-frequency and 5 common SNV associations detected in the 1x (19% of total) are not seen associated below that threshold in the imputed GWAS. As expected, there are significantly fewer (3.8%, P=0.01, one-sided chi-square proportion test) true associations in the imputed GWAS not recapitulated by the 1x study.

**Figure 5:**
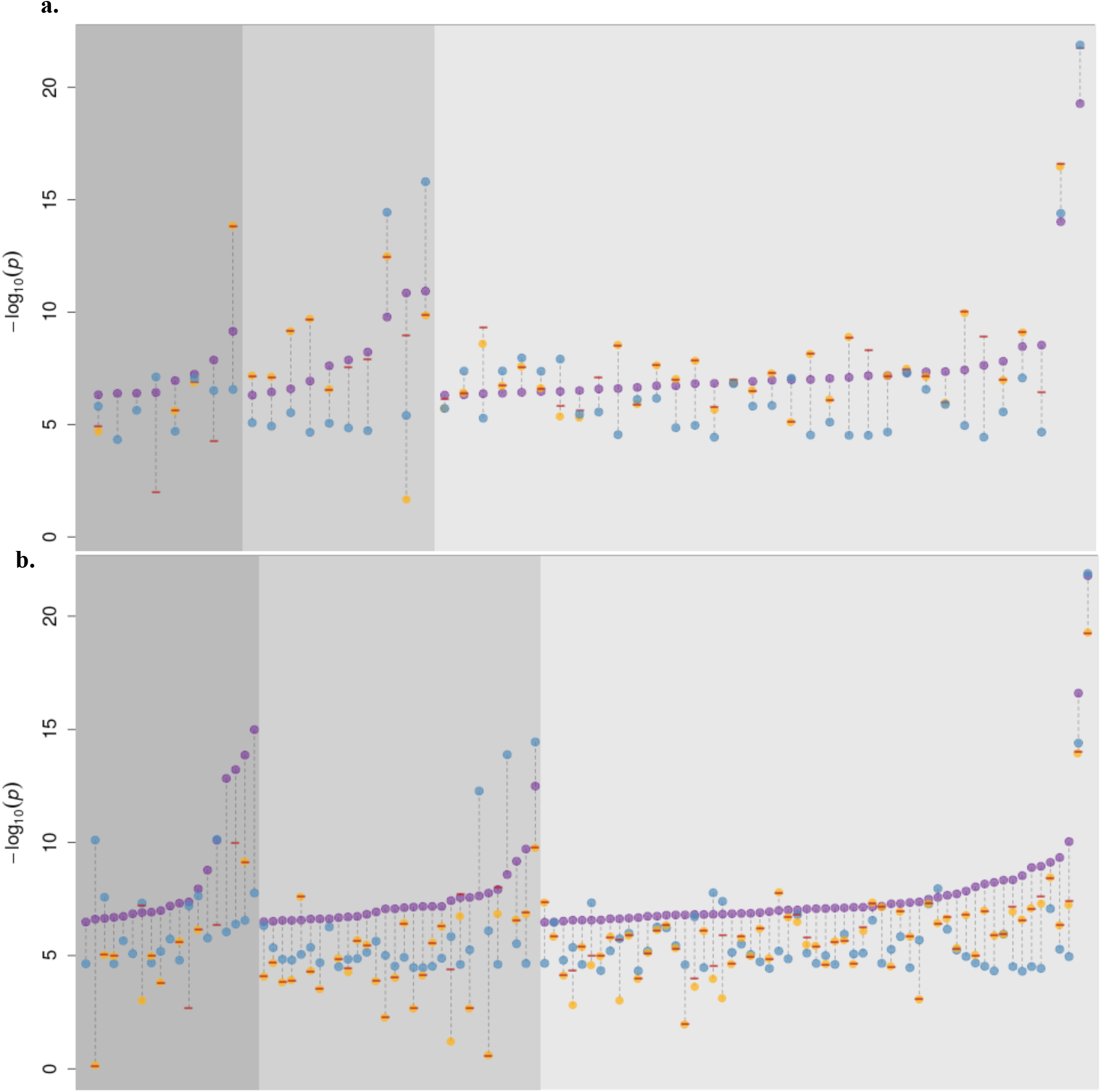
Association signals in the 1x WGS and imputed GWAS at p<5xl0^−7^ for 57 quantitative traits in 1,225 samples. Purple dots represent significant results in the 1x WGS (a.) and imputed GWAS (b.) analysis. Orange dots, if present, denote the p-value of the same SNP in the other study. Blue dots represent the association p-value in a larger (n=1,457) association study based on 22x WGS. Signals with a 22x WGS p-value above 5x10^−5^ were considered as false positives in both studies and excluded from the plot. Red dashes indicate the minimum p-value among all tagging SNVs in the other dataset (r^2^>0.8). Absence of an orange dot and/or a red dash means that the variant was not present and/or no tagging variant could be found for that signal in the other study.

## Discussion

In this work, we empirically demonstrate the relative merits of very low depth WGS both in terms of variant discovery and association study power for complex quantitative traits compared to GWAS approaches. However, the advantages of 1x WGS have to be weighed against compute and financial cost considerations. As of summer 2018, 1x WGS on the HiSeq 4000 platform was approximately half of the cost of a dense GWAS array (e.g. Illumina Infinium Omni 2.5Exome-8 array), the same cost as a sparser chip such as the Illumina HumanCoreExome array, and half of the cost of WES at 50x depth. By comparison, 30x WGS was 23 or 15 times more costly depending on the sequencing platform (Illumina HiSeq 4000 or HiSeqX, respectively). The number of variants called by 1x WGS is lower than high-depth WGS, but is in the same order of magnitude, suggesting comparable disk storage requirements for variant calls. However, storage of the reads required an average 650Mb per sample for CRAMs, and 1.3Gb per sample for BAMs.

Genome-wide refinement and imputation of very low depth WGS generates close to 50 times more variants than a GWAS chip. The complexity of the imputation and phasing algorithms used in this study is linear in the number of markers, linear in the number of target samples and quadratic in the number of reference samples (Browning and Browning, 2016), which results in a 50-fold increase in total processing time compared to an imputed GWAS study of equal sample size. In MANOLIS the genome was divided in 13,276 chunks containing equal number of SNVs, which took an average of 31 hours each to refine and impute. The total processing time was 47 core-years (see Methods and Supplementary Figure 2). This parallelisation allowed processing the 1,239 MANOLIS samples in under a month, and as imputation software continue to grow more efficient (Bycroft, et al., 2017), future pipelines should greatly simplify postprocessing of very low depth sequencing data.

As a proof of principle, we used imputed GWAS, 1x and 22x WGS in overlapping samples from an isolated population to assess how genotyping quality influences power in association studies. As we only wanted to study the implications of varying genotype qualities afforded by different designs on association p-values in a discovery setting, we considered only suggestively associated signals and did not seek replication in a larger cohorts for the discovered signals. In our study of 57 quantitative traits, we show that an 1x-based design allows the discovery of twice as many of the signals suggestively associated in the more accurate 22x WGS study, compared to the imputed GWAS design. Almost 10% of the suggestive signals arising in the 1x data are not discoverable in the imputed GWAS, but the great majority (96%) of imputed GWAS signals is found using the 1x.

The 1x-based study seems to discover more signals than the imputed GWAS across the MAF spectrum, and this remains true whether or not the signals are filtered for suggestive association p-value in the more accurate 22x based study (Supplementary Table 2). At first glance this suggests 1x WGS has better detection power than the imputed GWAS across the MAF spectrum, however it is unlikely that this is true for common variants, which are reliably imputed using chip data. Instead, this phenomenon is likely due to a slightly less accurate imputation than in the GWAS dataset caused by a noisier raw genotype input (Supplementary Text). This effect is marginal, as evidenced by genome-wide concordance measures (Figure 2) which are very high at the common end of the MAF spectrum. However, it is important to note that this slightly less accurate imputation can attenuate some signals as well as boosting others. For this reason, we would recommend relaxing the discovery significance threshold in 1x studies in order to capture those less well imputed, signal-harbouring variants, followed by rigorous replication in larger cohorts and direct validation of genotypes.

Our study’s intent was to focus on the performance on commonly used general-purpose tools for low-depth sequencing data in isolates, both for genotype calling (GATK) and imputation (BEAGLE, IMPUTE). There are ongoing efforts to leverage the specificities of both low-depth sequencing (Davies, et al., 2016)(https://www.gencove.com) and of isolated populations (Livne, et al., 2015). The popularity and long-term support of established generic methods is an advantage when running complex study designs, as has been shown in other isolate studies (Herzig, et al., 2018).

We show that very low depth whole-genome sequencing allows the accurate assessment of most common and low-frequency variants captured by imputed GWAS designs and achieves denser coverage of the low-frequency and rare end of the allelic spectrum, albeit at an increased computational cost. This allows very low depth sequencing studies to recapitulate signals discovered by imputed chip-based efforts, and to discover significantly associated variants missed by GWAS imputation (Gilly, et al., 2016). Although cohort-wide high-depth WGS remains the gold standard for the study of rare and low-frequency variation, very low-depth WGS designs using population-specific haplotypes for imputation remain a viable alternative when studying populations poorly represented in existing large reference panels.

## Methods

### Cohort details

The HELIC (Hellenic Isolated Cohorts; www.helic.org) MANOLIS (Minoan Isolates) collection focuses on Anogia and surrounding Mylopotamos villages on the Greek island of Crete. All individuals were required to have at least one parent from the Mylopotamos area to enter the study. Recruitment was primarily carried out at the village medical centres. The study includes biological sample collection for DNA extraction and lab-based blood measurements, and interview-based questionnaire filling. The phenotypes collected include anthropometric and biometric measurements, clinical evaluation data, biochemical and haematological profiles, self-reported medical history, demographic, socioeconomic and lifestyle information. The study was approved by the Harokopio University Bioethics Committee and informed consent was obtained from every participant.

### Sequencing

Sequencing and mapping for the 990 MANOLIS samples at 1x depth has been described previously (Gilly, et al., 2016), as well as for 249 MANOLIS samples at 4x (Southam, et al., 2017), and for 1,457 samples at 22x (Gilly, et al., 2018). For comparison, 5 samples from the cohort were also whole-exome sequenced at an average depth of 75x. We use a standard read alignment and variant calling pipeline using samtools(Li, et al., 2009) and GATK(McKenna, et al., 2010), which is described in detail in the Supplementary Text.

### Variant filtering

Variant quality score recalibration was performed using GATK VQSR v.3.1.1. However, using the default parameters for the VQSR mixture model yields poor filtering, with a Ti/Tv ratio dropoff at 83% percent sensitivity and a Ti/Tv ratio of 1.8 for high-quality tranches (Supplementary Figure 3.a). We therefore ran exploratory runs of VQSR across a range of values for the model parameters, using the dropoff point of the transition/transversion (Ti/Tv) ratio below 2.0 as an indicator of good fit (Supplementary Figure 4). A small number of configurations outperformed all others, which allowed us to select an optimal set of parameters. For the chosen set of parameters, false positive rate is estimated at 10%±5% (Supplementary Figure 3.b). Indels were excluded from the dataset out of concerns for genotype quality. We found that the version of VQSR, as well as the annotations used to train the model, had a strong influence on the quality of the recalibration (Supplementary Figure 4 and Supplementary Text).

### Comparison with downsampled whole genomes

For quality control purposes, reads from 17 of the well-characterised Platinum Genomes sequenced by Illumina at 50x depth (McCarthy, et al., 2016), and downsampled to 1x depth using samtools (Christopoulos, 1997) were included in the merged BAM file. VQSR-filtered calls were then compared to the high-confidence call sets made available by Illumina for those samples. 524,331 of the 4,348,092 non-monomorphic variant sites were not present in the high-confidence calls, whereas 1,246,403 of the 5,070,164 non-monomorphic high-confidence were not recapitulated in the 1x data. This corresponds to an estimated false positive rate of 12% and false negative rate of 24.6%. Both unique sets had a much higher proportion of singletons (corresponding to MAF < 2.9%) than the entire sets (57.9% vs 19.9% of singletons among 1x calls and 51% vs 18.1% among high-confidence calls), which suggests that a large fraction of the erroneous sites lies in the low-frequency and rare part of the allelic spectrum. However, genotype accuracy is poor, to the point where it obscures peculiarities in the distribution of allele counts (Supplementary Figure 5). Due to these being present in the 1000 genomes reference panel, we remove the 17 Platinum Genomes prior to imputation.

### Genotype refinement and imputation

#### Evaluation of pipelines

The authors of SHAPEIT (Delaneau, et al., 2013) advise to phase whole chromosome when performing pre-phasing in order to preserve downstream imputation quality. This approach is computationally intractable for the 1x datasets, where the smallest chromosomes contain almost 7 times more variants than the largest chromosomes in a GWAS dataset.

For benchmarking purposes, we designed 13 genotype refinement pipelines involving Beagle v4.0 (Browning and Browning, 2007) and SHAPEIT2 (Delaneau, et al., 2013) using a 1000 Genomes phase 1 reference panel, which we evaluated against minor allele concordance. All pipelines were run using the vr-runner scripts (https://github.com/VertebrateResequencing/vr-runner). Pipelines involving Beagle with the use of a reference panel ranked consistently better (Supplementary Figure 1), with a single run of reference-based refinement using Beagle outperforming all other runs. IMPUTE2 performed worst on its own, whether with or without reference panel; in fact the addition of a reference panel did not improve genotype quality massively. Phasing with Beagle without an imputation panel improved genotype quality, before or after IMPUTE2.

Halving the number of SNVs per refinement chunk (including 500 flanking positions) from the 4,000 recommended by the vr pipelines resulted in only a modest loss of genotype quality in the rare part of the allelic spectrum (Supplementary Figure 7), while allowing for a twofold increase in refinement speed. Genotype quality dropped noticeably for rare variants when imputation was turned on (Supplementary Figure 7), but remained high for low-frequency and common ones. A reference-free run of Beagle allowed to phase all positions and remove genotype missingness with no major impact on quality and a low computational cost. We also tested thunderVCF (Pollin, et al., 2008) for phasing sites, however, the program took more than 2 days to run on 5,000 SNV chunks and was abandoned.

#### Production pipeline for the MANOLIS cohort

For production, we used a reference panel composed of 10,244 haplotypes from the 1000 Genomes Project Phase 1 (n=1,092), UK10K (UK10K Consortium, et al., 2015) TwinsUK (Moayyeri, et al., 2013) and ALSPAC (Golding, et al., 2001) (n=3,781, 7x WGS), and 249 MANOLIS samples sequenced at 4x depth, which has been described before (Southam, et al., 2017). Alleles in the reference panel were matched to the reference allele in the called dataset. Positions where the alleles differed between the called and reference datasets were removed from both sources. Indels were filtered out due to poor calling quality.

The pipeline with best minor allele concordance across the board used Beagle v.4 (Browning and Browning, 2007) to perform a first round of imputation-based genotype refinement on 1,239 HELIC MANOLIS variant callsets, using the aforementioned reference panel. This was followed by a second round of reference-free imputation, using the same software.

#### Variant-level QC

Beagle provides two position level imputation metrics, allelic R-squared (AR2) and dosage R-squared (DR2). Both measures are highly correlated (Supplementary Figure 8.a). Values between 0.3 and 0.8 are typically used for filtering (Browning, 2014). In both 1x datasets 59% and 91% of imputed variants lie below those two thresholds, respectively. The distribution of scores does not provide an obvious filtering threshold (Supplementary Figure 8.b) due to its concavity. Since most imputed variants are rare and R-squared measures are highly correlated with MAF, filtering by AR2 and DR2 would be similar to imposing a MAF threshold (Supplementary Figure 8.c and d.). Moreover, due to a technical limitation of the vr-runner pipelines, imputation quality measures were not available for refined positions at the time of analysis, only imputed ones. Therefore, we did not apply any prior filter in downstream analyses.

#### Sample QC

Due to the sparseness of the 1x datasets, sample-level QC was performed after imputation. 5 samples were excluded from the MANOLIS 1x cohort following PCA-based ethnicity checks.

#### Comparison with WES

A set of high confidence genotypes was generated for the 5 exomes in MANOLIS using filters for variant quality (QUAL>200), call rate (AN=10, 100%) and depth (250x). These filters were derived from the respective distributions of quality metrics (Supplementary Figure 9).

When compared to 5 whole-exome sequences from each cohort, imputed 1x calls recapitulated 77.2% of non-monomorphic, high-quality exome sequencing calls. Concordance was high, with only 3.5% of the overlapping positions exhibiting some form of allelic mismatch. When restricting the analysis to singletons, 9105 (58%) of the 15,626 high-quality singletons in the 10 exomes were captured, with 21% of the captured positions exhibiting false positive genotypes (AC>1). To assess false positive call rate, we extracted 1x variants falling within the 71,627 regions targeted by the Agilent design file for WES in overlapping samples, and compared them to those present in the unfiltered WES dataset. 103,717 variants were called in these regions from WES sequences, compared to 58,666 non-monomorphic positions in the 1x calls. 1,419 (2.4%) of these positions were unique to the 1x dataset, indicating a low false-positive rate in exonic regions post-imputation.

**Genetic relatedness matrix**

Relatedness was present at high levels in our cohort, with 99.5% of samples having at least one close relative (estimated 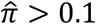) and an average number of close relatives of 7.8. In order to correct for this close kinship typical of isolated cohorts, we calculated a genetic relatedness matrix using GEMMA (Zhou and Stephens, 2012). Given the isolated nature of the population and the specificities of the sequencing dataset, we used different variant sets to calculate kinship coefficients. Using the unfiltered 1x variant dataset produced the lowest coefficients (Figure 10.a), whereas well-behaved set of common SNVs (Arthur, et al., 2017) produced the highest, with an average difference of 3.67x10^−3^. Filtering for MAF lowered the inferred kinship coefficients. Generally, the more a variant set was sparse and enriched in common variants, the higher the coefficients were. However, these differences only had a marginal impact on association statistics, as evidenced by a lambda median statistic difference of 0.02 between the two most extreme estimates of relatedness when used for a genome-wide association of triglycerides (Supplementary Figure 10.b). For our association study, we used LD-pruned 1x variants filtered for MAF<1% and Hardy Weinberg equilibrium p<1x10^−5^ to calculate the relatedness matrix, which translated into 2,848,245 variants for MANOLIS.

### Single-point association

#### Pipeline

For association, fifty-seven phenotypes were prepared, with full details of the trait transformation, filters and exclusions described in Supplementary Table 3. The ‘transformPhenotype’ (https://github.com/wtsi-team144/transformPhenotype) R script was used to apply a standardised preparation for all phenotypes. Association analysis was performed on each cohort separately using the linear mixed model implemented in GEMMA (Zhou and Stephens, 2012) on all variants with minor allele count (MAC) greater than 2 (14,948,665 out of 30,483,158 variants in MANOLIS). We used the aforementioned centered kinship matrix. GC-corrected p-values from the likelihood ratio test (p_lrt) are reported. Singletons and doubletons are removed due to overall low minor allele concordance. Signals were extracted using the peakplotter software (https://github.com/wtsi-team144/peakplotter) using a window size of 1Mb.

#### Data Access

The HELIC genotype and WGS datasets have been deposited to the European Genome-phenome Archive (https://www.ebi.ac.uk/ega/home): EGAD00010000518; EGAD00010000522; EGAD00010000610; EGAD00001001636, EGAD00001001637. The peakplotter software is available at https://github.com/wtsi-team144/peakplotter, the transformPhenotype app can be downloaded at https://github.com/wtsi-team144/transformPhenotype.

## Acknowledgements

The authors thank the residents of the Mylopotamos villages for taking part in the study. The MANOLIS study is dedicated to the memory of Manolis Giannakakis, 1978–2010. This study makes use of data generated by the UK10K Consortium, derived from samples from the TwinsUK Cohort and the Avon Longitudinal Study of Parents and Children (ALSPAC). A full list of the investigators who contributed to the generation of the data is available from www.UK10K.org, last accessed April 29, 2016. The GATK program was made available through the generosity of Medical and Population Genetics program at the Broad Institute, Inc. This research has been conducted using the UK Biobank Resource using Application Number 13745. This work was funded by the Wellcome Trust [098051] and the European Research Council [ERC-2011-StG 280559-SEPI]. Funding for UK10K was provided by the Wellcome Trust under award WT091310. The authors thank Sophie Hackinger and Bram Prins for proofreading the article. The authors also thank Heather Elding, William Astle, Tao Jiang Adam Butterworth and Nicole Soranzo for their contributions to a previous version of the manuscript.

